# Transcriptional bursts explain autosomal random monoallelic expression and affect allelic imbalance

**DOI:** 10.1101/649285

**Authors:** Anton J.M. Larsson, Björn Reinius, Tina Jacob, Tim Dalessandri, Gert-Jan Hendriks, Maria Kasper, Rickard Sandberg

## Abstract

Transcriptional bursts render substantial biological noise in cellular transcriptomes. Here, we investigated the theoretical extent of monoallelic expression resulting from transcriptional bursting and how it compared to the amounts of monoallelic expression of autosomal genes observed in single-cell RNA-sequencing (scRNA-seq) data. We found that transcriptional bursting can explain frequent observations of autosomal monoallelic gene expression in cells. Importantly, the burst frequency largely determined the fraction of cells with monoallelic expression, whereas both burst frequency and size contributed to allelic imbalance. Allelic observations deviate from the expected when analysed across heterogeneous groups of cells, suggesting that allelic modelling can provide an unbiased assessment of heterogeneity within cells. Finally, large numbers of cells are required for analyses of allelic imbalance to avoid confounding observations from transcriptional bursting. Altogether, our results shed light on the implications of transcriptional burst kinetics on allelic expression patterns and phenotypic variation between cells.

## Introduction

Cells of identical background and type may show phenotypic differences (***Symmons and Raj, 2016***) and stochasticity in the transcriptional process itself is known to generate cellular variability (***Elowitz et al., 2002***). In mammalian cells, diploidy may give rise to phenotypic differences due to the unequal expression of two functionally different alleles. Indeed, since mammalian genes are transcribed in discrete bursts (***Elowitz et al., 2002***; ***Chubb et al., 2006***; ***Raj et al., 2006***; ***Suter et al., 2011***) there are periodic fluctuations in the abundance of transcripts originating from each allele, which have implications for the allelic expression within cells over time and may further extend to phenotypic differences (***Reinius and Sandberg, 2015***).

The introduction of allele-sensitive single-cell RNA-sequencing revealed that RNA from substantial numbers of autosomal genes were only detectable from a single allele in individual cells at any given time point (***Deng et al., 2014***). This autosomal random monoallelic expression (aRME) is consistent with the concept of transcriptional bursting (***Nicolas et al., 2017***), in particular since subsequent work demonstrated that the allelic patterns were primarily due to a stochastic process in somatic cells rather than a mitotically heritable characteristic (***Reinius et al., 2016***). Furthermore, allele-specific RNA FISH of autosomal genes *in situ* has shown that transcriptional bursting can explain the observed aRME in three selected cases (***Symmons et al., 2019***). However, the explicit relationship between aRME and transcriptional burst kinetics has not been systematically explored.

The two-state model of transcription (***Peccoud and Ycart, 1995***; ***Raj et al., 2006***) (***Figure 1A***) has been extensively used to investigate quantitative relationships between burst kinetics and other gene-level measurements (***Raj et al., 2006***; ***Suter et al., 2011***; ***Kim and Marioni, 2013***). This model has four gene-specific parameters that may accommodate different kinds of kinetics, mainly characterized by the burst frequency and size, with frequency normalized by mRNA degradation rates. A main limitation to broadly investigating the implications of transcriptional burst kinetics has been the challenge of obtaining reliable allelic estimates of transcriptional burst kinetics in diploid cells for sufficiently large numbers of genes. However, this barrier was recently overcome by advances in the inference of transcriptional burst kinetics from allele-sensitive scRNA-seq (***Kim and Marioni, 2013***; ***Larsson et al., 2019***; ***Jiang et al., 2017***), culminating in the demonstration that enhancers drive burst frequencies and that core promoter elements effect burst size (***Larsson et al., 2019***).

**Figure 1.**
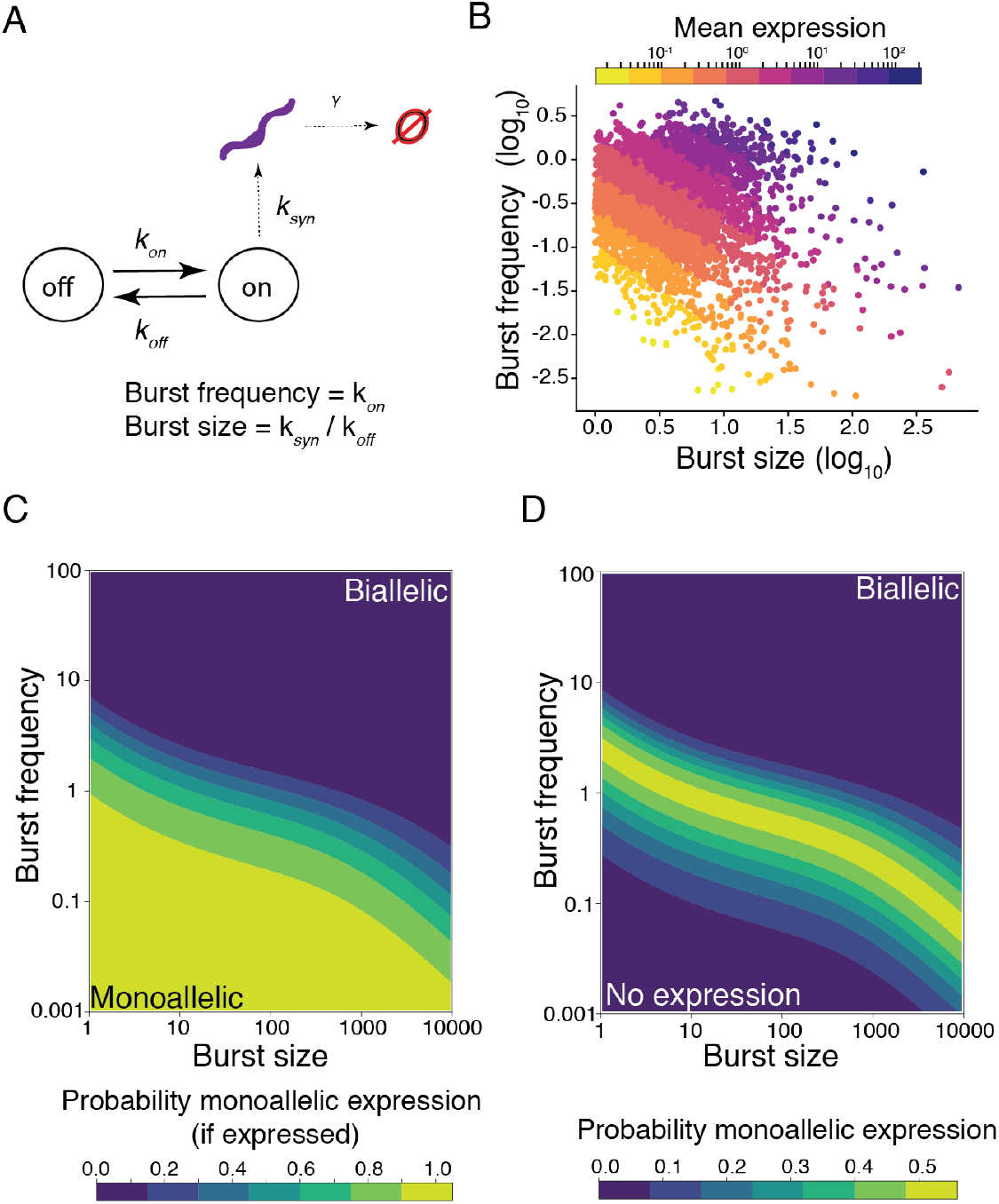
The theoretical effect of transcriptional bursting on dynamic random monoallelic expression. **(A)** Illustration of the model used for transcriptional burst kinetics. The time for the gene to transition are given by the exponentially distributed parameters *k*_*on*_ (from off to on) and *k*_*off*_ (from on to off). While the gene is active, the gene is transcribed at rate *k*_*syn*_. The burst frequency is given by *k*_*on*_ and the average number of transcripts produced in a burst (burst size) is given by *k*_*syn*_/*k*_*off*_. **(B)** A scatter plot showing burst frequency and burst size estimates from the C57 allele of autosomal genes in mouse fibroblasts (*n* = 4,905 genes), where each gene is colored based on the mean expression level of that gene (mean number of observed UMIs per cell). Figure based on data from (***Larsson et al., 2019***). **(C)** Contour plot of the conditional probability of observing monoallelic expression when there is expression of that gene in the parameter space of burst frequency and size. **(D)** Contour plot of the probability of observing monoallelic expression in the parameter space of burst frequency and size, irrespectively if the gene is expressed or not.

In the present study, we used parameters of burst kinetics inferred from thousands of genes of a mouse cross breed (CAST/EiJ × C57BL/6J) (***Larsson et al., 2019***). We showed that most genes have a pattern of expression that is consistent with the concept of dynamic aRME (***Reinius and Sandberg, 2015***), wherein monoallelic expression occurs due to independent bursts of transcription from each allele. The inferred kinetic parameters predict the fraction of monoallelic expression on each allele with high precision. We furthermore showed, *in vitro* and *in vivo*, that the fraction of monoallelic expression is mainly driven by the frequency of transcriptional bursts rather than burst sizes, whereas allelic imbalance is a consequence of both burst frequencies and size.

## Results

We first investigated the theoretical impact of transcriptional burst kinetics on random monoallelic gene expression, using the two-state model of transcription (***Figure 1A***). Importantly, a vast number of combinations of burst frequencies and sizes results in the same level of mean expression, which is readily observable in scRNA-seq data (***Figure 1B***). To investigate where in parameter space monoallelic gene expression patterns occur, we first calculated the probabilities of generating monoallelic expression for two alleles with identical transcriptional kinetics throughout kinetics parameter space. The parameters (*k*_*on*_, *k*_*off*_, *k*_*syn*_) describe the distribution of transcripts at steady state (**Methods**) and give the probability of detecting a transcript from the allele of a given gene, *P* (*n* > 0|*k*_*on*_, *k*_*off*_, *k*_*syn*_), where *n* is the number of transcripts. By conditioning the probabilities on the total probability of expression *P*(monoallelic|expressed), we find that genes with low burst frequency (*k*_*on*_) and size (*k*_*syn*_/*k*_*off*_) are always monoallelically expressed given that there is expression at all (***Figure 1C***). A combination of high burst frequency and size gives exclusively rise to biallelic states, while intermediate combinations of these extremes lie on a spectrum in-between. If we do not condition on expression, we can observed a ridge of states of monoallelic expression where biallelic and no expression dominate on either side of the ridge (***Figure 1D***).

As transcriptional burst kinetics are inferred on each allele independently (***Larsson et al., 2019***), we asked to what extent the measurements of monoallelic and biallelic expression from scRNA-seq experiments concur with the model predictions. To this end, we used inferred transcriptional burst kinetics of genes on the CAST and C57 alleles from 224 individual primary mouse fibroblasts (***Larsson et al. (2019***), CAST/EiJ × C57BL/6J) which resulted in 4,905 autosomal genes with reliable kinetics inferred for both alleles after filtering **(Methods)**. In these cells, we calculated the fraction of monoallelic expression from both alleles and considered whether the monoallelic expression was affected by the burst kinetics of the genes. We found that the observed fraction of monoallelic expression show the same relationship with the burst kinetics as predicted by theory, with a ridge of monoallelic expression appearing across the parameter space of burst kinetics (***Figure 2A***). Furthermore, we estimated the probabilities of observing a cell which is either silent, biallelic, monoallelic on CAST or monoallelic on C57 for all genes, assuming that transcription occurs independently on each allele. The predicted fractions of cells in each state were highly correlated with the observed fraction of cells in each category (Spearman *r* = 0.98, 0.99, 0.92 and 0.93 respectively), (***Figure 2B***) demonstrating that modelling transcription using the two-state model at each allele independently is in agreement with experimental allelic expression analyses by scRNA-seq.

**Figure 2.**
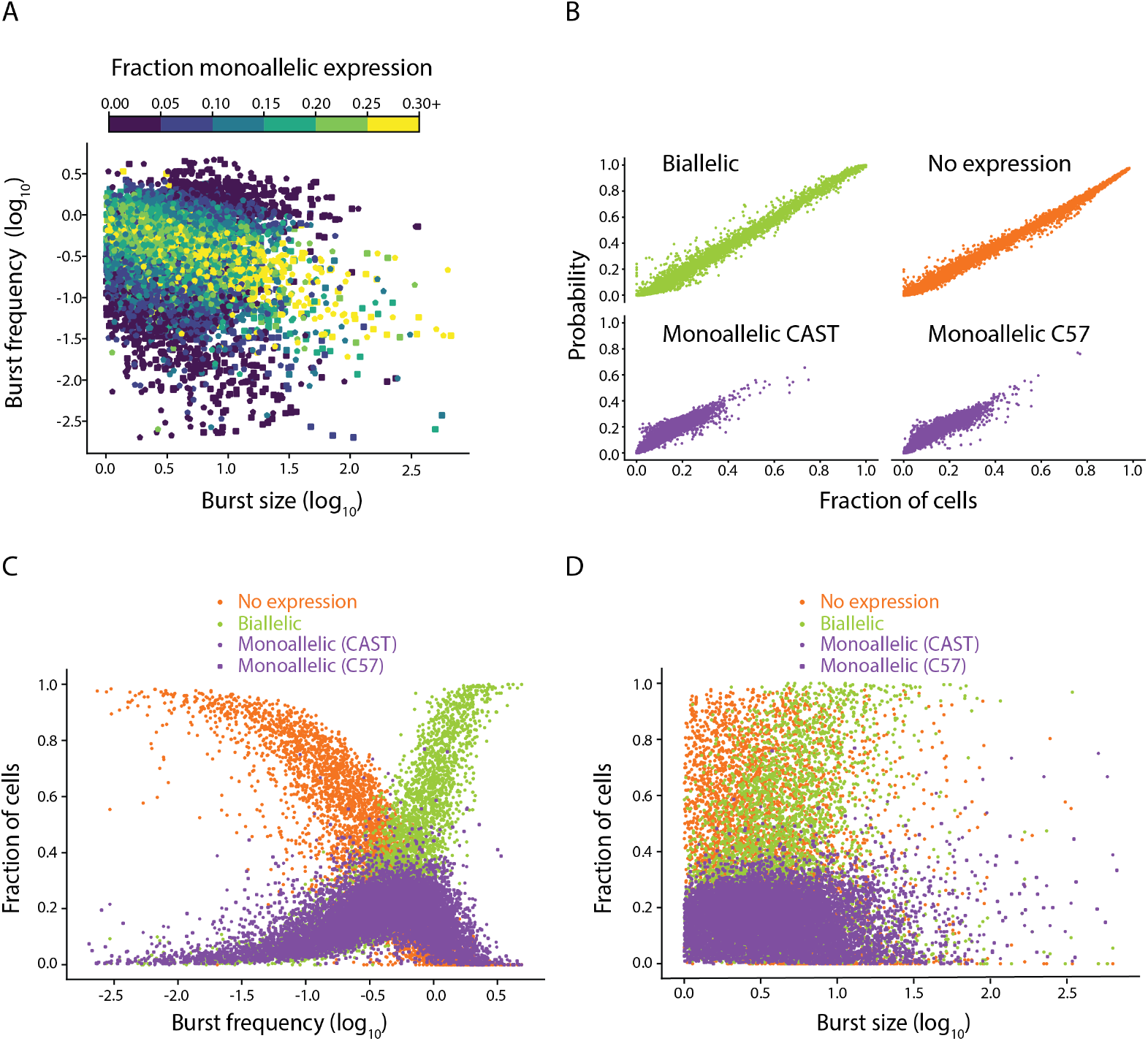
The relationship between transcriptional burst kinetics and dynamic random monoallelic expression in primary mouse fibroblasts. **(A)** A scatter plot showing burst frequency and burst size estimates from both alleles in mouse fibroblasts (C57 square, CAST pentagon, n=4,905 autosomal genes), where each gene is colored based on the fraction of cells which expressed the gene monoallelically from that allele (*n* = 224 cells). **(B)** Correlations between the predicted and observed fraction of cells with: no expression (orange), biallelic expression (green), monoallelic expression from the CAST allele and monoallelic expression from the C57 allele (violet), *n* = 4,905 genes. **(C-D)** The observed fraction of cells with biallelic, monoallelic (on CAST or C57) and no detectable expression for 4,905 autosomal genes with bursting kinetics inferred in mouse fibroblasts. Shown as a function of **(C)** burst frequency and **(D)** and burst size.

The theoretical results indicated that changes in aRME can be achieved by modulation of either burst frequency or size. To investigate which parameter of transcriptional bursting is the most decisive factor for aRME in the cell, we examined the profile of burst frequency and size in relation to monoallelic expression to isolate their relative contributions. The burst frequency of the CAST and C57 alleles compared to their fraction of monoallelic expression showed a striking relationship (***Figure 2C***). At lower burst frequencies, we observed very low amounts of monoallelic expression because there was predominantly no expression from either allele in any cell. The fraction of monoallelic expression increased as the burst frequency increased, up until the point where biallelic expression became the predominant observation and monoallelic expression then declined. This relationship was also clear in the theoretically predicted case which demonstrated that our model predictions were consistent with the biological data (***Supplementary Figure 1***). The same analysis on burst size showed that the distribution of monoallelic expression was almost uniform over burst size with a tendency of genes with large burst sizes to have more biallelic expression (***Figure 2D***). Therefore, while burst size has the theoretical capability to influence the amount of monoallelic expression in cells, it plays a minor role relative to burst frequency. The predominant role of burst frequencies in determining monoallelic gene expression can also be seen in the slope of the ridge of monoallelic expression (***Figure 1D***,***Figure 2A***)

To extend the inference and analyses of transcriptional burst kinetics to cell types *in vivo*, we sequenced individual cells from dorsal skin of the same mouse cross breed (C57BL/6JxCAST/EiJ). In the 354 single-cell transcriptomes (post quality control), we identified 10 clusters (***Figure 3A***) that could be assigned to cell types of variable heterogeneity using existing skin single-cell transcriptomics data (***Joost et al., 2016***). The relationship between burst kinetics and random monoallelic gene expression for cells *in vivo* was consistent with the data from primary fibroblasts (***Supplementary Figure 2***).

**Figure 3.**
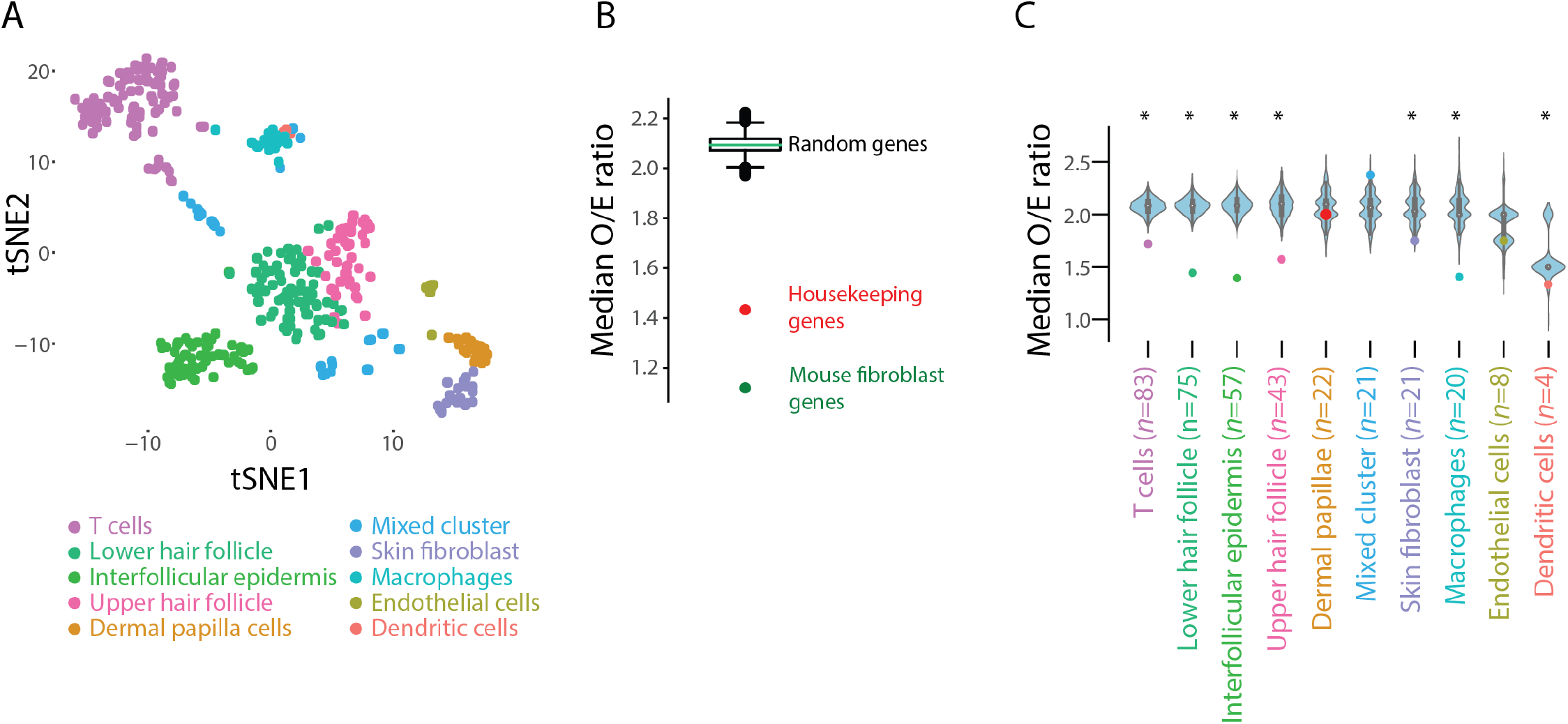
Heterogeneity in cell clusters from an *in vivo* experiment in mouse skin measured by observed-to-expected biallelic expression. **(A)** t-distributed stochastic neighbour embedding (tSNE) of the skin cells, colored by SNN-based clustering (*n* = 354 cells). **(B)** The median observed-to-expected (O/E) ratio of biallelic expression, comparing the theoretical predictions from burst kinetics to that observed in all cells without stratifying cells to clusters. Boxplot show median O/E biallelic expression from random sets of genes (*n* = 3,727 autosomal genes and 100,000 permutations) whereas the red dot show the O/E ratio when analyzing housekeeping genes in all cells. For comparison, the analyses of all genes in primary fibroblasts are shown in green. **(C)** The median O/E ratio of biallelic expression within cell clusters shown as colored dots. These were compared to randomly selected cells of the same size (*n* = 83, 75, 57, 43, 22, 21, 21, 20, 8, 4 cells respectively, 1,000 permutations for each cluster). Asterisk denotes significance at *α* = 0.05.

Due to the heterogeneous cellular composition of certain clusters, we could quantify how well each cell-type cluster predicted its own biallelic expression based on the model of independent allelic transcriptional bursting (**Methods**). We anticipated that high heterogeneity within a cell cluster would show an underestimation of predicted biallelic expression due to subsets of heterogeneous cells with higher burst frequency for certain genes. Indeed, the median observed-to-expected ratio of biallelic expression (O/E ratio) based on all cells (irrespective of clustering) indicated a clear transcriptome-wide underestimation of biallelic expression (median = 2.1, *n* = 10,543 genes).

To examine the potential of allelic-expression modelling as an unbiased method to assess the degree of heterogeneity within groups of cells, we first examined housekeeping genes as they would have less cell-type-specific transcriptional burst regulation compared to other genes and thereby have observed biallelic call frequencies closer to the expected value (an O/E ratio closer to 1). Indeed, housekeeping genes had a significantly lower O/E ratio compared to randomly selected subsets of genes, and were close to the ratio observed in the fibroblast cells (*p* < 10^−5^, permutation test, ***Figure 3B***). Importantly, the stratification of cells into clusters greatly improved the O/E ratio compared to randomly selected sets of cells, with the exception of three clusters (containing mixed unassigned cells, partly endothelial and dermal papillae cells, *p* < 10^−3^, permutation test, ***Figure 3C***). Therefore, the observed-to-expected biallelic expression due to transcriptional bursting may be a valuable metric to quantify the heterogeneity within an assigned cell cluster.

Investigating gene expression at the discrete level of monoallelic and biallelic expression is natural at the single-cell level as it is frequently observed due to transcriptional bursting. However it is also important to assess the consequences of transcriptional bursts on the continuum of allelic bias or imbalance. To this end, we calculated the theoretical probabilities of the allele of a gene occurring in larger or equal amounts than the other allele, *P* (*C*57 > *CAST*), *P* (*CAST* > *C*57) and *P* (*CAST* = *C*57), in the primary fibroblasts. The probability of equal expression is dominated by the outcome of no expression on either allele, which is predictably related to the burst frequencies of the two alleles of the gene (***Supplementary Figure 3***). Most genes have very similar kinetics between the two alleles and therefore a close to equal probability of unequal expression for each allele, as measured by *P* (*C*57 > *CAST*|*C*57 ≠ *CAST*) (***Supplementary Figure 3***).

The probabilities were in good agreement with the observed fractions of allelic bias (***Figure 4A***). By comparing the fold changes in burst size and frequency between alleles to their observed fraction of allelic bias, we found that the relative differences in transcriptional burst kinetics in burst frequency as well as size tended to affect allelic bias for that gene (***Figure 4B***). Interestingly, simultaneous relative changes in both burst frequency and size may cancel each other out. For example, a reduction in burst size may be compensated by an increase in burst frequency (***Figure 4B***; visualized along the diagonal of the scatter plot). By using linear regression with allelic bias as the dependent variable, we determined that relative changes in both burst frequency and size together explain the allelic bias to a high degree (*R*^2^ = 83.7%) and both relative changes have significant impact on allelic bias (***Table 1***). Thus, the imbalanced gene expression of CAST and C57 alleles arise from genetic variation in regulatory enhancers and promoter sequences (***Larsson et al., 2019***), affecting both burst frequencies and sizes.

**Table 1.**
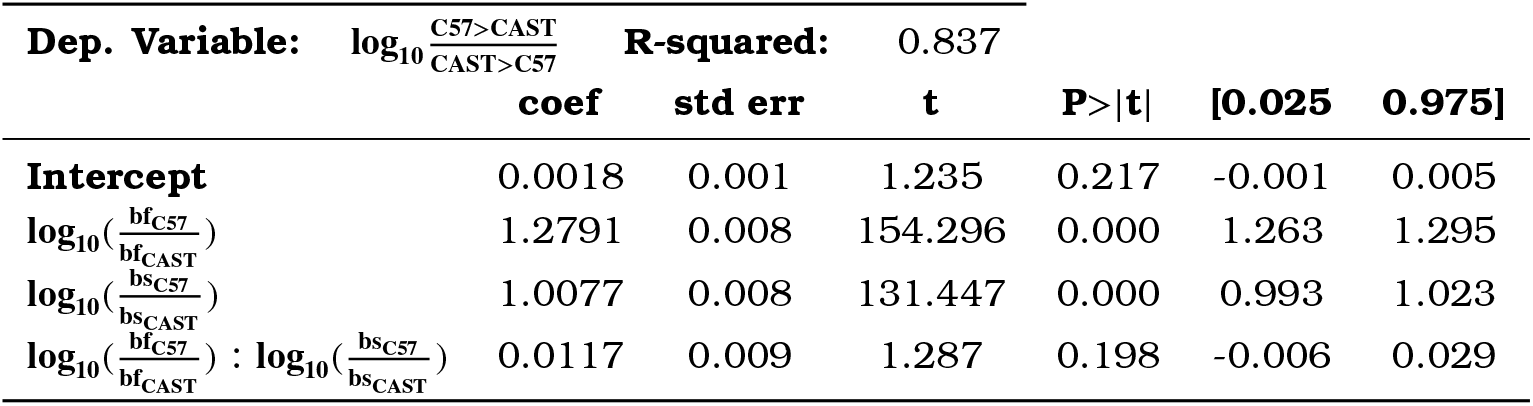
Ordinary least squares regression results for the effect of burst kinetics on allelic imbalance. bf: burst frequency, bs: burst size, coef: linear regression coefficient, std err: standard error, t: t-statistic.

**Figure 4.**
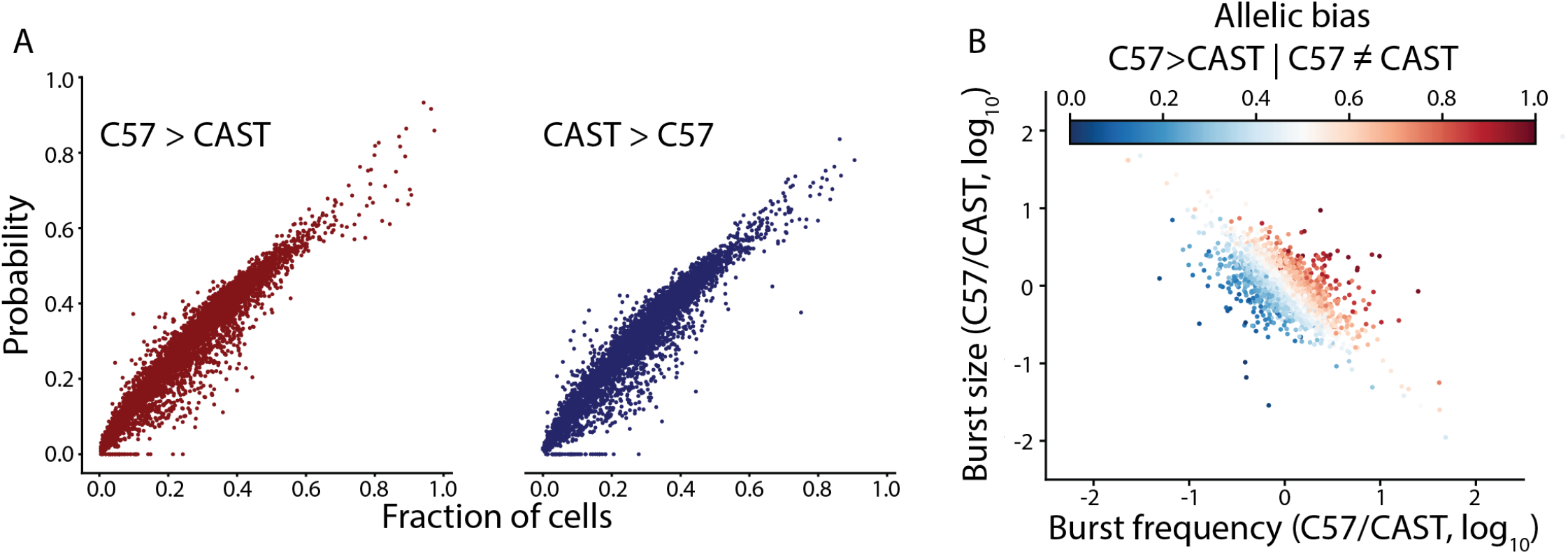
Allelic bias is affected by relative changes in both burst frequency and size. **(A)** Comparison between the probability of observing allelic imbalance between the alleles and the actual fraction of cells with the imbalance (*n* = 4,905 autosomal genes). **(B)** The relative allelic differences in burst kinetics for each gene, colored by their allelic bias (*n* = 4,905 genes).

To determine the extent to which transcriptional bursting may give rise to false positives in studies of allelic imbalance in single cells, we simulated the expression from two alleles with kinetics identical to those inferred from the C57 allele for different number of cells (***Figure 5A***, *n* = 10, 20, 50, 100, 1,000 and 10,000 cells). We then estimated the allelic imbalance for all genes based on a model that expected equal expression from both alleles. At a low number of sequenced cells, the variance in expression due to transcriptional bursting severely impacts the allelic imbalance measurements and give rise to a high number of false positives, but becomes increasingly stable with a higher number of cells (***Figure 5B***). In relation to mean expression, we find that it is only for low-expressed genes that false positive allelic imbalance becomes frequent, and this declines as the number of simulated cells increases (***Figure 5C***).

**Figure 5.**
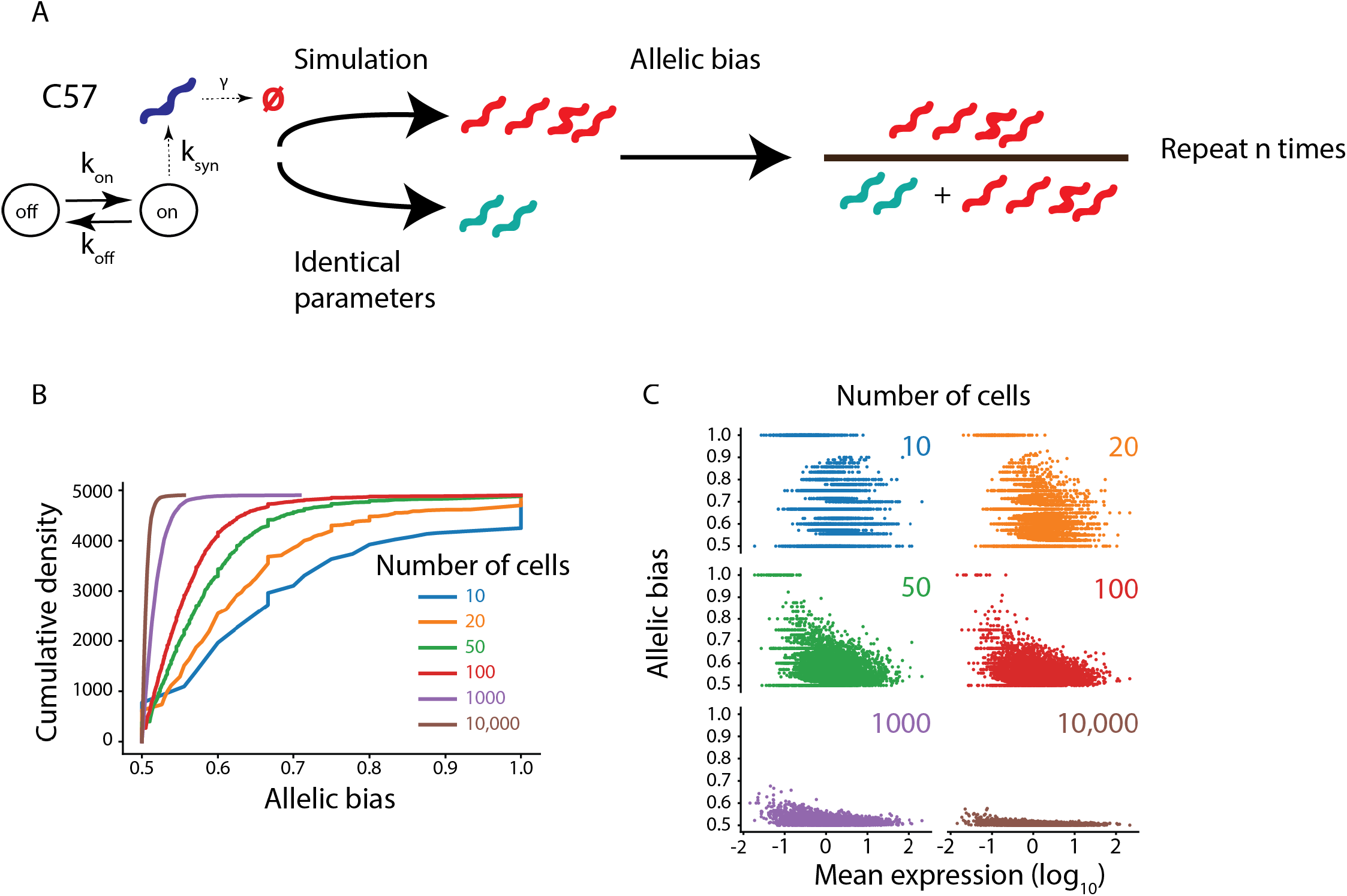
Low-expressed genes frequently show false positive allelic imbalance due to transcriptional bursting. **(A)** Outline of the simulation strategy. **(B)** The cumulative distribution of allelic bias of the simulated genes with the same kinetics (*n* = 4,905 autosomal genes), where allele with the highest allelic bias is the chosen value for each gene. **(C)** The relationship between the mean expression of a gene and allelic bias based on the number of simulated cells (*n* = 4,905 genes).

## Discussion

In this study, we explored if transcriptional burst kinetics can explain observed patterns of random monoallelic gene expression of autosomal genes (***Reinius and Sandberg, 2015***). We report a striking resemblance between theory and observations in allelic expression patterns, which provides explicit support that transcriptional burst kinetics result in frequent monoallelic expression of autosomal genes. Moreover, we conclude that burst frequency largely determines the extent to which a gene is monoallelically expressed in somatic diploid cells, whereas both burst frequency and size contribute to allelic imbalance.

We additionally explored to what extent transcriptional bursts can lead to spurious observations of allelic imbalance in cell populations. In full agreement with single-molecule RNA FISH analyses of allelic gene expression in cells *in vivo*, we also found that lowly expressed genes can be falsely identified as having allelic imbalance simply due to their stochastic transcription (***Symmons et al., 2019***). The number of false positives of allelic imbalance that we present is likely a conservative estimate, as we do not include genes for which the burst kinetic inference procedure fails and therefore exclude many genes which have even lower expression level than the genes analyzed herein. Here, we also did not extensively model the technical noise present in the library preparation or the variability during sequencing. However, simply increasing the numbers of cells investigated for example in an RNA-sequencing experiment might not fully mitigate this effect, since cellular proportions of the cell type constituents combined with their cell-type specific expression might still result in a number of genes which are subject to noisy allelic imbalance estimates due to transcriptional bursting. To what extent this can explain the scarce numbers of autosomal genes with clonally propagated monoallelic gene expression is currently unknown, but it is interesting in this context to note that most previously identified genes were detected at very low levels (around two RNA transcripts per cell on average when expressed, (***Reinius and Sandberg, 2015***; ***Eckersley-Maslin et al., 2014***; ***Gendrel et al., 2014***). It is clear that future studies of monoallelic gene expression and allelic imbalance in diploid cells need to consider the consequences of stochastic transcription.

There is current great excitement in using single-cell RNA-sequencing to identify and characterize cell types, sub-types and cellular states throughout human tissues and in model organisms (***Regev et al., 2017***; ***Han et al., 2018***). In these efforts, methods that can determine the quality of cell type assignments are important and allelic modelling of transcriptional kinetics could be a strategy for unbiased evaluation of the homogeneity within assigned cell clusters. Here, we suggest biallelic status as a measure of cellular heterogeneity within a cluster of cells. We show that housekeeping genes and our *in vivo* cell clusters from mouse skin show an observed-to-expected biallelic expression ratio which is lower than what would be expected by chance. If there is heterogeneity with regards to the kinetic bursting parameters between the cells we would expect to observe biallelic expression at a much higher frequency than what would be predicted and therefore a higher biallelic O/E ratio. In model organisms, this can easily be achieved by profiling cells of F1 offspring of characterized parent strains. However, phased haplotypes may also be obtained for human cells in the transcribed parts of the genome from the single-cell RNA-sequencing data itself (***Edsgärd et al., 2016***), which would allow for such modelling.

Transcriptional bursting results in considerable cellular heterogeneity in cases of two functionally different alleles (e.g. see ***Montag et al. (2018)***). It is however not clear if such variation has phenotypic consequences. Interestingly, transcriptional bursting was recently shown to impact T-cell linage commitment (***Ng et al., 2018***) which raise the intriguing question whether cell fate decisions in general could be affected by stochastic transcription. Since burst frequency is preferentially encoded in enhancer regions (***Larsson et al., 2019***) it is likely that mutations in *cis*-regulatory sequences or trans-activating factors may affect the penetrance of phenotypes. This may be particularly relevant in the case of lineage commitment, which exhibits switch-like irreversible activation. In the case of allelic imbalance, burst size may play a relatively larger role in terms of phenotype. The relative abundances of protein products resulting from translation of the two different alleles may be affected by burst size which could be relevant in the case of phenotypes that result due to the stoichiometric constraints present in signalling pathways and gene networks.

## Methods and Materials

### The two-state (beta-poisson) model

The model used for stochastic gene expression is a particular case of a birth-and-death process in a Markovian environment. In short, the model has the states (*i, n*) where *i* ∊ {0, 1} indicates if the gene is active or not, and *n* is the number of RNA transcripts in the cell.

In the off state, the gene can turn on with the rate *k*_*on*_. In the on state, the gene can turn off with rate *k*_*off*_ and produce one RNA transcript with the rate *k*_*syn*_. Regardless of the state, one RNA transcript can be degraded with rate *γ*. At the steady state of this process, the stationary distribution can be shown to be described by the Poisson-beta distribution, in which we let

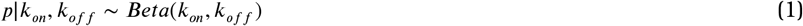

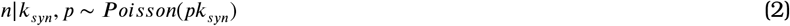

The resulting marginal distribution *P* (*n*|*k*_*on*_, *k*_*off*_, *k*_*syn*_) is the probability distribution for the amount of RNA transcripts observed at steady state given the rates *k*_*on*_, *k*_*off*_ and *k*_*syn*_.

### Inference of transcriptional burst kinetics

We inferred kinetic bursting parameters for 7,191 and 7,186 genes for the C57 and CAST allele respectively from 224 F1 cross-breed (CASTxC57) adult tail fibroblasts. We then applied a goodness-of-fit test to these parameters to assess how well the parameters describe the data and found that 5,586 and 5,601 genes fit the model for the C57 and CAST alleles respectively. The intersection of kinetic parameters between both alleles after accounting for the goodness-of-fit test and removing genes on the X chromosome result in 4,905 usable genes for our analysis. The method to infer these parameters given allele-sensitive scRNA-seq data and the goodness-of-fit test is described in ***Larsson et al. (2019***) and the code for doing so is available at https://github.com/sandberg-lab/txburst.

### Calculating the probabilities and observed fractions for silent, biallelic, monoallelic expression and allelic bias

Given the parameters inferred in Larsson et. al. we can calculate the probability of allele expressing a given gene or not at the time of sampling. We define a function of the probability of observing *k* UMI counts for an allele of gene *g* given the parameters, *P* (*K* = *k*|*k*_*on*_, *k*_*off*_, *k*_*syn*_).

With the resulting genes we can calculate the probabilities of an allele of the genes not being observed, i.e.

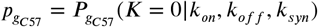

and

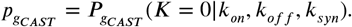

This allows us to calculate the probabilities of:

- Probability of expression on no allele 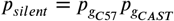
- Probability of monoallelic expression on the C57 allele 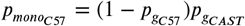
- Probability of monoallelic expression on the CAST allele 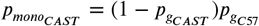
- Probability of biallelic expression 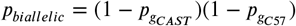

These probabilities assume that the alleles burst independently. Since our probabilities closely follow the observed fractions this assumption seems to be correct. The contour plots in ***Figure 1C*** and ***Figure 1D*** are based on 100×100 parameter combinations where *k*_*off*_ is varied to change burst size while is held constant at 100.

For each gene, we calculated the fraction of no expression, monoallelic on C57, monoallelic on CAST and biallelic expression by averaging the following conditional statements over the cells where *n*_*allele*_ refers to the number of actual UMI counts for that allele in that cell:

- No expression: *n*_*C*57_ = 0 and *n*_*CAST*_ = 0
- Monoallelic expression on the C57 allele: *n*_*C*57_ > 0 and *n*_*CAST*_ = 0
- Monoallelic expression on the CAST allele: *n*_*CAST*_ > 0 and *n*_*C*57_ = 0
- Biallelic expression: *n*_*CAST*_ > 0 and *n*_*C*57_ > 0

For the comparisons between predicted and observed values we used spearman correlations.

We then calculated the theoretical probabilities of the allele of a gene occurring in larger amounts than the other allele by considering the probability of

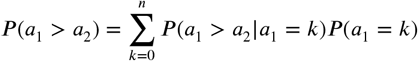

where *a*_1_ and *a*_2_ is the number of RNA transcripts from allele 1 and 2 respectively and *n* is the highest number of RNA transcripts for *a*_1_ with a non-zero probability of being observed. For each gene we then find three probabilities *P*(*C*57 > *CAST*), *P*(*CAST* > *C*57) and *P*(*CAST* = *C*57)

### Preparation, sequencing and analysis of skin cells

Skin tissue was dissected from 9 week old female F1 offspring of matings between CAST/EiJ and C57BL/6J mice (approval by the Swedish Board of Agriculture, Jordbruksverket: N95/15). Cells were dissociated from skin as described in Joost et al. 2016, or using GentleMACS (Miltenyi Biotec); with both methods giving similar cell yields and viability. Briefly, for the GentleMACS method, dorsal skin was cut and minced into small pieces (approximately 1×1mm) and incubated in HBSS (Sigma) + 0.04% BSA (Sigma) + 0.2% Collagenase Ia (Sigma) at 37°C for 60 minutes with occasional agitation. Thereafter this slurry was processed on a GentleMACS (Miltenyi Biotec) with 2x Program D, cell-strained (70um) and washed. Residual tissue was further treated with HBSS + 0.05% Trypsin-EDTA (Sigma) at 37°C for 15 minutes and processed likewise. For the Joost et al method, GentleMACS dissociation was substituted with manual disaggregation by smashing tissue fragments against a cell strainer with the piston from a 5 mL syringe. Cells were sorted into 384-well plate by FACS, and subject to Smart-seq2 single-cell RNA-sequencing library creation (***Picelli et al., 2014***). The single-cell libraries were sequenced on an Illumina HiSeq4000, the sequence fragments aligned to the mouse genome (mm10) and summarized into expression levels (RPKMs) and allele-resolved expression, as previously described (***Larsson et al., 2019***). Single-cell data was processed and analysed using Seurat (version 2.3.4), including log-normalization, regression of the total number of detected reads, identification of genes with most biological variation (n=1000), SNN-based clustering (distances in PCA-space, using the 20 top principal components), followed by manual curation of certain clusters (endothelial cells, dermal papillae, dendritic cells, mixed cluster). Cells with less than 100k mapped reads were excluded from the analysis (30 cells). The allelic expression levels were used for transcriptional burst kinetics inference, as described above. The discrepancy in scale between burst size values in the smart-seq2 data and the ***Larsson et al. (2019)*** data is due to UMIs, for a more detailed discussion see ***Larsson et al. (2019)***.

### Assessing heterogeneity of cell-type clusters by observed-to-expected biallelic expression

We sequenced 384 cells from the dorsal skin (in telogen) of the C57BL/6JxCAST/EiJ cross breed using smart-seq2 (***Picelli et al., 2014***), see above for details. To calculate the observed-to-expected ratio of biallelic expression for each gene, we calculated the expected fraction of cells with biallelic expression based on the model of independent bursts of transcription,

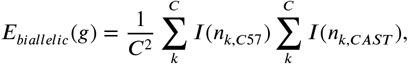

where *C* is the number of cells, *k* the *k*th cell and *I*(*n*) is the indicator function

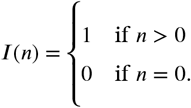

We then calculate the observed number of cells with biallelic expression for that gene, *O*_*biallelic*_(*g*), to combine them to obtain *O*_*biallelic*_(*g*)/*E*_*biallelic*_(*g*). The list of housekeeping genes was obtained from (***Li et al., 2017***).

### Ordinary least squares regression of the effect of burst kinetics on allelic bias

We used the OLS module of the statsmodels package in Python with the formula:

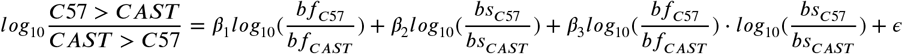

where bf is burst frequency and bs is burst size.

### Calculating allelic bias based on simulated observations

For ***Figure 5*** we used the burst kinetics parameters inferred from the C57 allele and simulated observations for each gene twice for a varying number of observations (*n* = 10, 20, 50, 100, 1000, 10000 observations). We then calculated max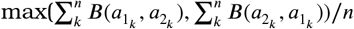 for each gene where 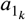 and 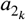 are the observed values for the *k*th simulated pair of observations and

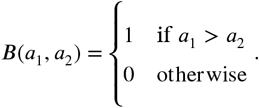

## Data Availability

The single-cell RNA-seq data generated in this study has been deposited at Gene Expression Omnibus (ID pending). The computational workflows will be uploaded as Jupyter notebooks to our Github account.

## Acknowledgments

We thank the members of the Sandberg laboratory for their input to this project. This work was supported by grants to R.S. from the European Research Council (648842), the Swedish Research Council (2017-01062), the Knut and Alice Wallenberg’s foundation (2017.0110), the Bert L. and N. Kuggie Vallee Foundation and the Goran Gustafsson Foundation.

## Author Contributions

Conceived the study: R.S. Analyzed data: A.J.M.L. and T.J. Interpreted data: A.J.M.L., B.R. and R.S. Collected and prepared single-cell libraries of mouse dorsal skin cells: G.J.H. and T.D. Wrote the manuscript: R.S and A.J.M.L. with input from B.R. Supervised the study: M.K. and R.S.

---

**Supplementary Figure 1.**
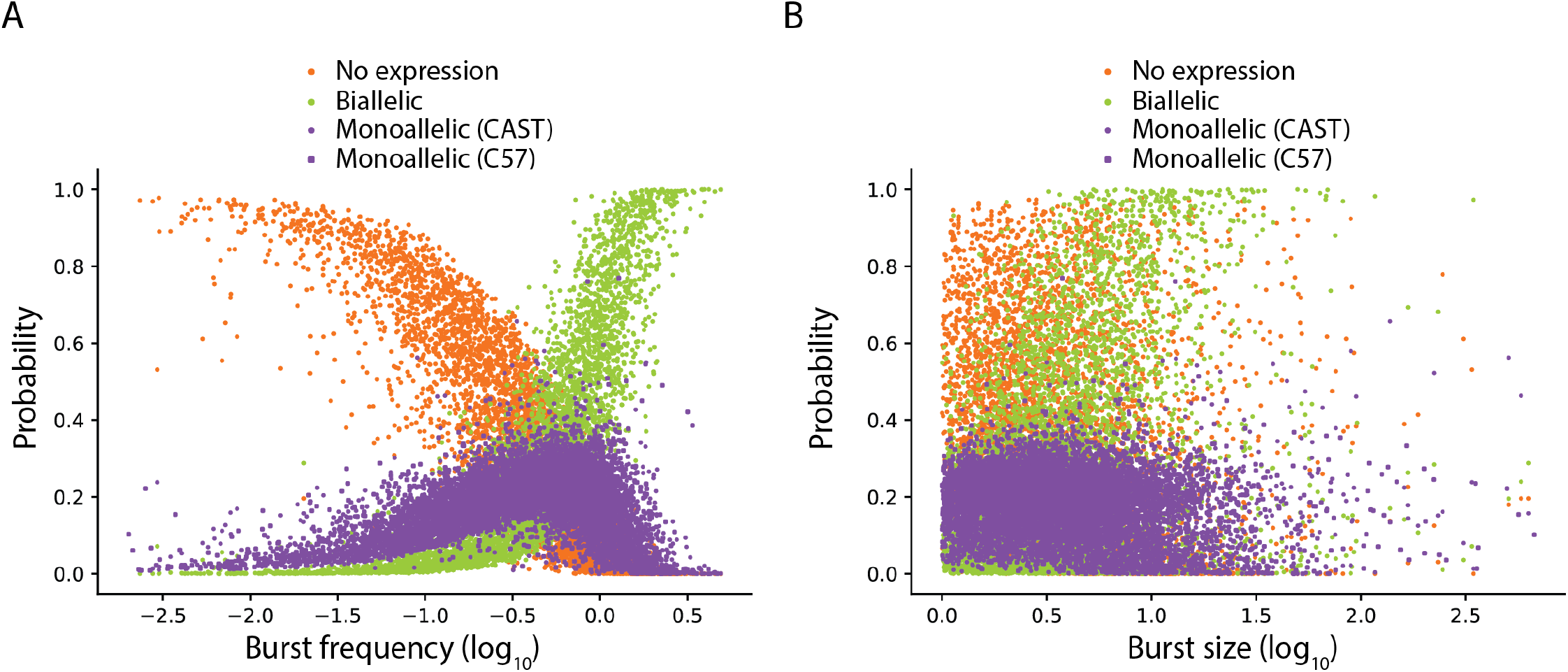
The probabilities of having biallelic, monoallelic (from CAST or C57) or no detectable expression directly predicted from the two-state model of transcription, as a function of burst frequency **(A)** and burst size **(B)**. Note, the results are almost identical to the observed dependencies in single-cell RNA-seq data (shown in **Figure 2C**,**Figure 2D**).

**Supplementary Figure 2.**
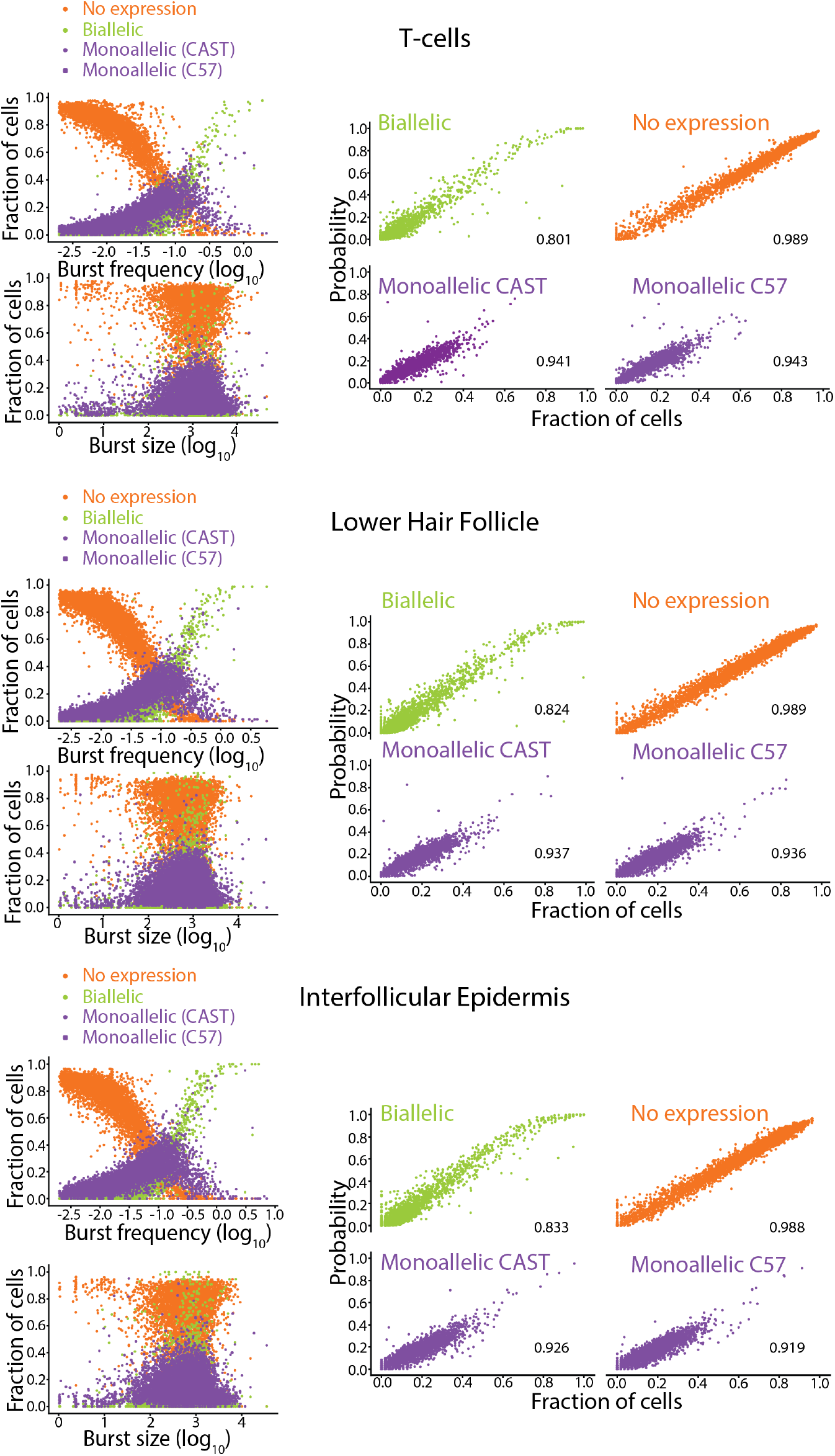
Same analysis as in **Figure 2** for the three largest cell type clusters in the mouse skin *in vivo* data. **Top:** T-cells (*n* = 4299 genes from 83 cells), **Middle:** Lower Hair Follicle cells (*n* = 5807 genes from 75 cells), **Bottom:** Interfollicular Epidermal cells (*n* = 5145 genes from 57 cells). **Right:** Relationship between inferred burst kinetics and allelic expression patterns. **Left:** Correlations between predicted and actual allelic expression patters, spearman correlation coefficient in the bottom right.

**Supplementary Figure 3.**
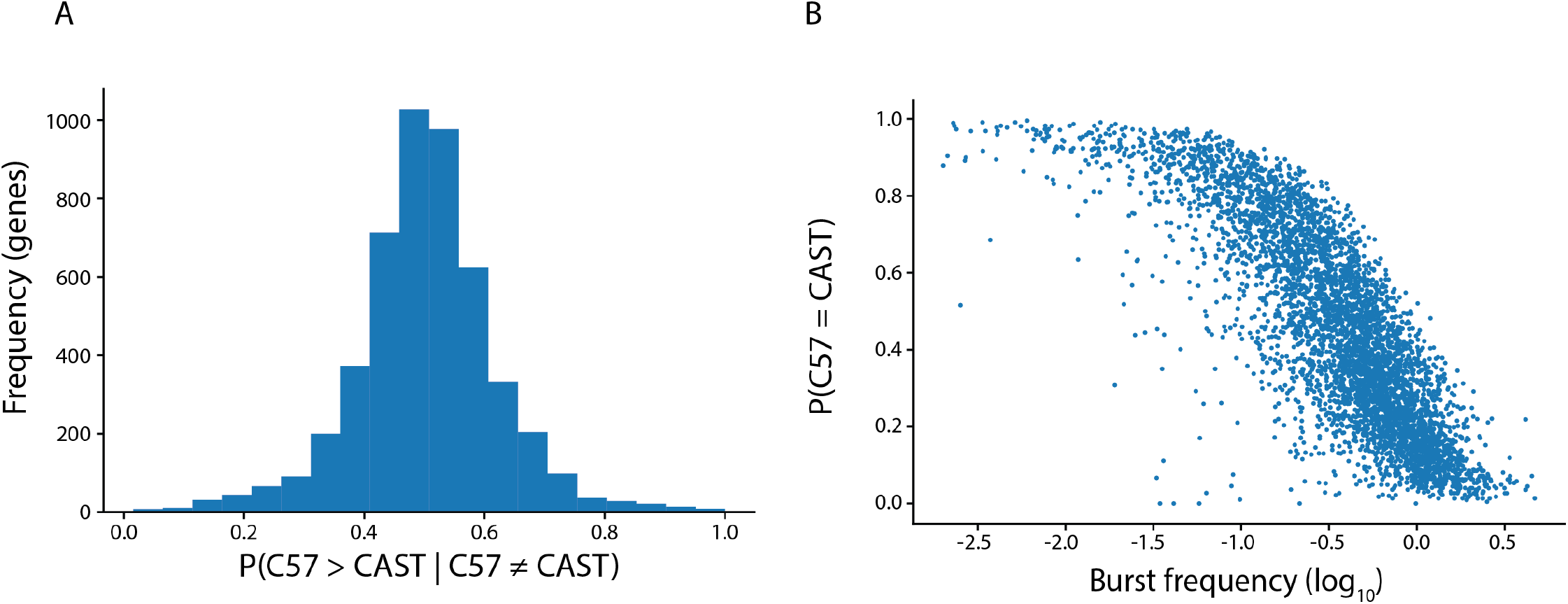
**(A)** Distribution of *P*(*C*57 > *CAST*|*C*57 ≠ *CAST*) (*n* = 4905 genes). **(B)** Relationship between burst frequency and equal expression (which is dominated by no expression on either allele).

